# Assessing PDMS Biocompatibility in Microfluidic Applications: Toxicity and Survival Outcomes in *C. elegans*

**DOI:** 10.64898/2025.12.11.693574

**Authors:** Kin Gomez, Kirill Efimenko, Jan Genzer, Adriana San-Miguel

## Abstract

Polydimethylsiloxane (PDMS), often assumed to be biocompatible, is widely used in microfluidic devices and biomedical research. Here, we systematically assess the organismal effects of PDMS network components and their leachates using *Caenorhabditis elegans* as a whole-animal model. We demonstrate that uncrosslinked vinyl-terminated PDMS (v-PDMS) chains, which comprise the majority of a PDMS network and are known to diffuse into aqueous environments, exhibit acute, environmentally-dependent toxicity. Low-molecular-weight v-PDMS (6 kDa) caused mild lethality in nutrient-rich S-Medium but significantly higher mortality in minimal S-Buffer, showing that media composition strongly influences toxic effects. Adding cholesterol, calcium, or magnesium notably reduced v-PDMS-induced lethality, whereas trace metals increased it. Using a DAF-16::GFP reporter strain, we show that cholesterol influences organismal stress responses to v-PDMS exposures. Progeny from starved parents showed full resistance to v-PDMS, suggesting transgenerational stress memory plays a role in reducing PDMS toxicity. We also find that linear siloxanes cause modest but significant lethality, whereas cyclic siloxanes do not. The PDMS crosslinker TDSS, however, provides partial protection when present with v-PDMS, revealing diverse biological effects among PDMS network precursors. Overall, these results show that PDMS-derived components are not universally harmless and that susceptibility depends greatly on environmental conditions, sterol levels, and physiological history. Our findings emphasize the importance of carefully evaluating PDMS formulations for biomedical use and offer a framework for assessing polymer leachate toxicity in living organisms.

## 1. Background/Introduction

Biomaterials play a crucial role in the fabrication of medical devices, drug delivery systems, and microfluidics, where biocompatibility is a key concern. Polydimethylsiloxane (PDMS) is a widely used silicone-based polymer with unique chemical and physical properties that enable diverse uses. PDMS oils are also used as lubricants^1^, water barriers^2^, and in cosmetic products such as topical creams and shampoos^3^. PDMS can also be found in food, due to its anti-foaming properties^3^. PDMS can be cured into a solid network by using a crosslinking agent, often in the presence of a catalyst. These PDMS networks result in a transparent soft elastomer widely considered to be biocompatible, that is resistant to many solvents, chemically inert, and thermally stable^4^. These characteristics expand the material*’*s applications to include coatings^4,5^, medical implants^6,7^, lenses^8^, and drug delivery systems^9,10^. PDMS is also the preferred material for fabricating microfluidic devices^3,11^.

Although PDMS elastomers are considered to be chemically stable formulations, it is well-documented that unreacted PDMS components can leach out^12–14^ under certain conditions. Breast implants have been reported to *“*bleed*”*, meaning that short molecular weight silicone molecules leach out of the network into the patient*’*s tissues^7,14^. In microfluidic devices, leaching has been shown to influence cell culture outcomes, raising concerns about its potential biological effects ^13,15^. While PDMS is widely regarded as biocompatible, recent studies suggest that certain components in PDMS formulations can interact with biological molecules, modifying proteins and altering metabolic functions in studied organisms, including barnacle larvae and mammalian cells^13,15–19^. Despite its widespread use in the medical, cosmetic, and food industries, the biological effects of PDMS exposure remain poorly understood, particularly at the organismal level. Little is known about the impact of PDMS and its components on simple model organisms, which limits our understanding of the broader ecological and physiological consequences of their use.

To address this knowledge gap, we investigated the potential biological interactions of PDMS components with *Caenorhabditis elegans* (*C. elegans*), a widely used model organism in toxicology and developmental biology^20–22^. *C. elegans* is particularly well-suited to assess biomaterial toxicity due to its short lifecycle, transparency, and well-characterized metabolic and stress response pathways^23^. The organism*’*s ability to respond to environmental stressors such as temperature, oxidants, xenobiotics, and osmotic challenges makes it an ideal system for evaluating how unreacted PDMS components affect development, behavior, and survival^23,24^. Additionally, *C. elegans* is frequently studied within PDMS-based microfluidic devices, making it particularly relevant for studying PDMS interactions^25–27^. Here, we investigate the interactions between *C. elegans* and PDMS components leached into the culture medium. Our results reveal that these interactions are formulation-dependent and vary across PDMS components, with some exhibiting toxic effects while others appear protective. Moreover, we find that the composition of the culturing medium significantly modulates these outcomes, with nutrient-rich environments either mitigating or exacerbating the observed toxicity.

## 2. Results

### 2.1. Toxicity of Sylgard 184 formulation

To assess whether unreacted components from a PDMS network leach into aqueous solutions at levels that might be toxic, we exposed *C. elegans* to aqueous leachates from cured Sylgard 184 at three different base-to-curing agent (A:B) ratios: 5:1, 10:1, and 20:1. Sylgard 184 is a silicone commercial formulation that contains 30-60 wt% of fumed silica to improve mechanical properties of cured elastomer films. The A:B=10:1 ratio is recommended by the manufacturer^28^. Henceforth, we refer to Sylgard 184 as PDMS and ignore the effect of fumed silica and other fillers and processing improvement components in Sylgard 184. The alternative formulations were selected for two reasons: (1) to simulate improperly mixed elastomers with a localized excess of one component, and (2) to reflect the common practice of modifying PDMS ratios to adjust mechanical properties, with higher curing agent content yielding stiffer features^29,30^ and higher base content yielding more flexible ones^31–33^. The cured elastomers were shredded to average sizes of 3.72 ± 0.08 mm, 3.68± 0.08 mm, 3.24 ± 0.08 mm for 5:1, 10:1, and 20:1 PDMS compositions respectively (Figure 1a), and incubated in a buffer solution suitable for handling worms (S-Buffer) at a ratio of 2 g of PDMS per 1 mL of buffer for 24 hours at room temperature. After incubation, the aqueous phase was filtered to remove any residual particulate matter. The resulting leachate solutions were then used as culture media for *C. elegans* L4 larvae, simulating the conditions the animals might encounter during exposure to PDMS-based microfluidic devices. Worm viability was assessed 24 hours post-exposure. Surprisingly, we found that L4 *C. elegans* larvae cultured in media pre-incubated with 5:1-cured PDMS, exhibited significant mortality. In contrast, no mortality was observed in worms exposed to leachates from PDMS cured at 10:1 or 20:1. In fact, survival was higher than in control conditions (Figure 1b).

**Figure 1.**
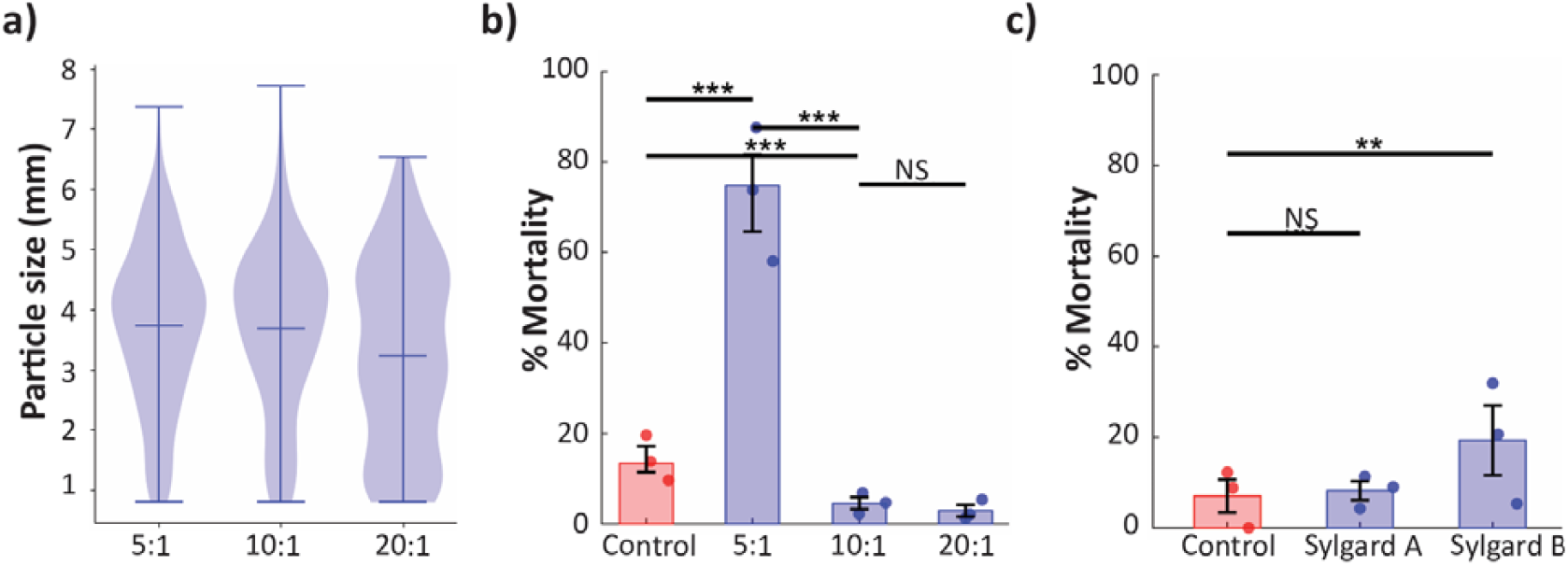
Mortality of *C. elegans* larvae exposed to Sylgard 184 leachates. (a) Cross-sectional area of shredded PDMS fragments (b) Leachates from cured PDMS prepared at different base:curing agent ratios (5:1, 10:1, 20:1). (b) Media equilibrated with Sylgard 184 Part A or Part B. Data shown as mean ± SEM, with weighted averages indicated by star symbols. Significance determined by Fisher*’*s exact test (p < 0.05, *p < 0.01, **p < 0.001). Approximately 100 worms per replicate per condition.

These results suggest that the toxic leachates observed in PDMS cured with excess curing agent originate from components present in Sylgard 184 component B, as only the condition with elevated proportions of component B resulted in increased worm mortality. Conversely, the improved survival observed in groups exposed to leachates from PDMS cured with excess elastomer base suggests that protective components responsible for this effect are likely present in Sylgard-184 Part A. To test whether the individual components of Sylgard 184 could induce similar effects to those of the leachates from cured elastomers, each component was mixed separately with the aqueous medium and incubated for 24 hours. The aqueous phase was then collected and used to culture *C. elegans* larvae. Consistent with the findings from the cured PDMS leachate experiments, exposure to media incubated with Sylgard 184 elastomer base (Part A) did not induce mortality in *C. elegans*, and survival rates were comparable to those of the control group. However, exposure to Part B resulted in a significant increase in lethality; unexpectedly, the effect was less pronounced than that observed with leachates from the 5:1-cured PDMS network (Figure 1c). These differences may indicate that specific components in the uncured mixtures become bioactive only upon network formation.

### 2.2. v-PDMS toxicity depends on molecular weight and media components

(α,ω-vinyl-terminated polydimethylsiloxane) v-PDMS chains (Figure 2a) constitute the main ingredient in PDMS formulations. In Sylgard 184, these chains make up more than 60% of Part A, and about 15–40% of the mass of Part B^34^, and have been shown to leach into culture media from microchannels^13^. To evaluate the potential bioactivity of v-PDMS leachates in *C. elegans*, we exposed L4-stage larvae to aqueous media (S-Medium) that had been equilibrated with v-PDMS of varying average molecular weights, following the same mixing preparation protocol described in the previous section. Initial results showed that exposure to low-molecular-weight v-PDMS (6 kDa) induced mild toxicity in worms cultured in S-Medium. In contrast, higher-molecular-weight variants produced no significant effects (Figure 2b).

**Figure 2.**
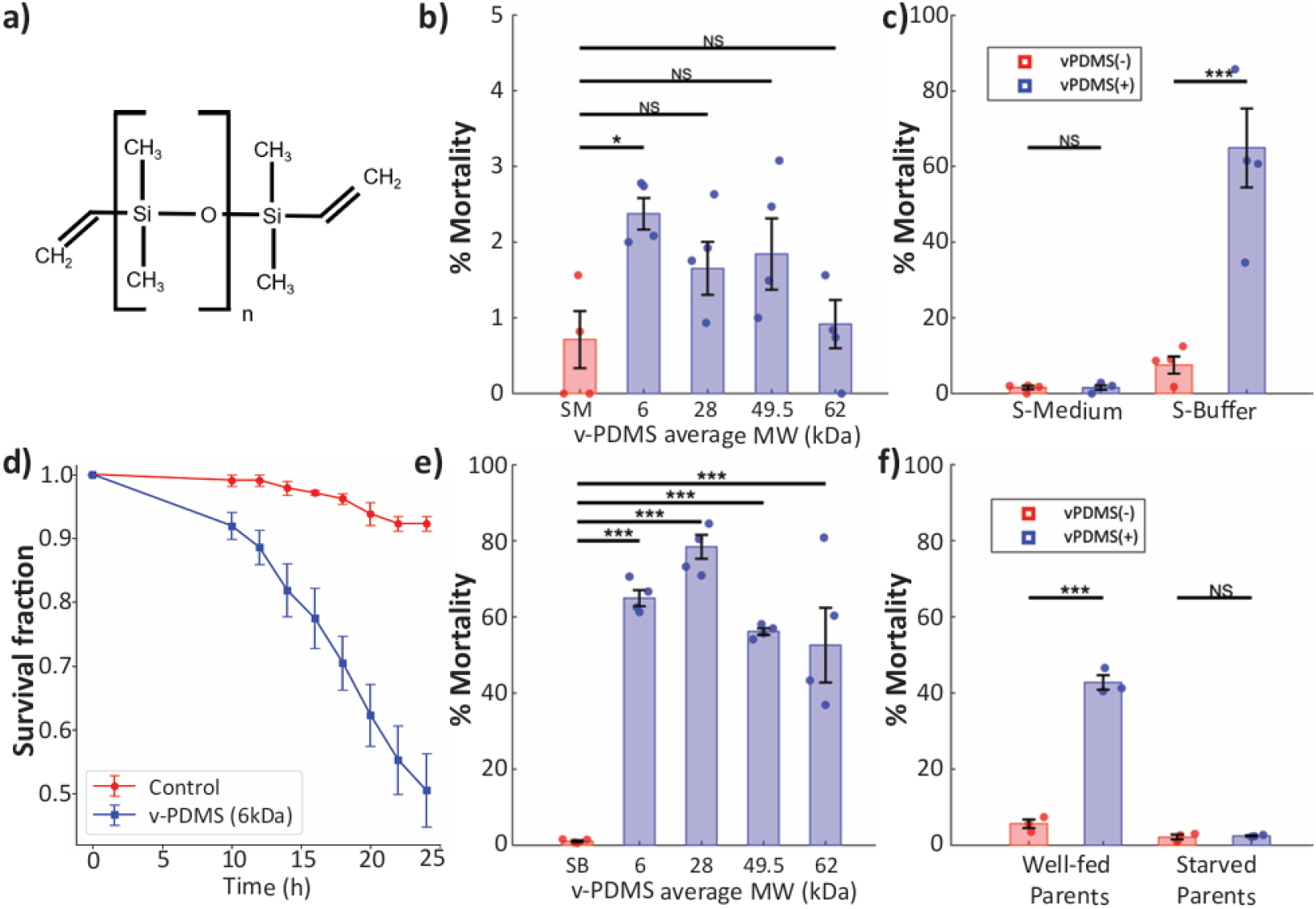
Survival of *C. elegans* exposed to v-PDMS oils under different conditions. (a) Chemical structure of (α,ω-vinyl-terminated polydimethylsiloxane) v-PDMS. (b) Mortality with v-PDMS of different molecular weights when cultured in S-Medium, significant only for 6 kDa chains. (c) Increased lethality of 6 kDa v-PDMS in S-Buffer compared to S-Medium. (d) Survival curves of L4 cultured in S-Buffer with and without v-PDMS. (e) All tested chain lengths induce mortality in S-Buffer. (f) Progeny of starved worms show full resistance to 6 kDa v-PDMS. Data as mean ± SEM, weighted averages indicated by stars. Fisher*’*s exact test with Bonferroni correction.

To determine whether components of the used aqueous media could contribute to v-PDMS toxicity, we compared S-Medium and S-Buffer as aqueous solutions. S-Buffer is a minimal medium typically used for short-term handling or stress assays that contains potassium phosphate buffer and NaCl. S-Medium contains S-Buffer in addition to cholesterol, potassium citrate, and other salts, as well as trace metals that support bacterial growth and worm development. We hypothesized that culturing worms under more nutritionally limited conditions and higher temperatures could heighten their sensitivity to v-PDMS. Increasing the temperature to 25°C did not significantly affect survival (Supplementary Figure 1). However, substituting S-Medium with S-Buffer resulted in a marked increase in v-PDMS-induced lethality (Figure 2c). Time-course assays further confirmed that 6 kDa v-PDMS causes a progressive decline in worm survival, with significant mortality observed within 24 hours of exposure (Figure 2d). Given the results of these assays, all experiments presented in this study were performed using S-Buffer as culture media unless specified otherwise. To further investigate the effect of v-PDMS molecular weight, we repeated the exposures using S-Buffer as the culture medium. Under these conditions, all tested v-PDMS molecular weights exhibited a substantially higher level of toxicity than experiments with S-media. However, a clear relationship between molecular weight and lethality was no longer observed (Figure 2e).

While performing these experiments, we observed that all worms derived from recently thawed populations survived exposure to v-PDMS in biological replicates. Given that *C. elegans* is known to exhibit transgenerational stress resistance following starvation^35,36^, we hypothesized that differences in developmental history could explain the differences in mortality between populations. To test this idea, we compared two populations: one that had been well-fed for five generations and another derived from a population that had experienced starvation during the L1 stage. We found that the progeny of starved worms exhibited complete resistance to v-PDMS toxicity (Figure 2f). This finding suggests that the harmful effects of v-PDMS can be attenuated through endogenous stress-resistance pathways that can be activated via transgenerational epigenetic inheritance. To ensure consistency, and based on this finding, all the other experiments presented here were conducted using worm populations that had been well-fed for at least five generations.

### 2.3. Calcium, magnesium, and cholesterol account for the media-dependent resistance to v-PDMS

To determine the cause of the increased toxicity observed in S-Buffer compared to S-Medium, we conducted toxicity experiments using modified variations of S-Buffer supplemented with different S-Medium components. S-Buffer serves as the base solution for S-Medium but lacks essential components necessary for the normal development of *C. elegans* and the growth of its bacterial food source, *E. coli* ^37^. To identify which components of S-Medium contribute to the reduced toxicity observed relative to S-Buffer, we supplemented S-Buffer individually with each of the major S-Medium additives (cholesterol, essential salts, and trace metals) and assessed worm survival following exposure to media incubated with 6 kDa v-PDMS using the same preparation and exposure protocol described previously. Since the base formulation for all liquid media used here is S-Buffer, all have similar pH (Supplementary Table 1). When the worms were cultured on S-Buffer supplemented with cholesterol, a significant decrease in toxicity was observed, suggesting that cholesterol plays a protective role against the toxic effects of v-PDMS (Figure 3a). However, the presence of trace metals in S-Medium had the opposite effect, enhancing the toxicity of v-PDMS leachates rather than mitigating it (Figure 3a). S-Medium contains magnesium sulfate and calcium chloride, essential minerals for *C. elegans* long-term viability^37^, and potassium citrate as a chelating agent and buffer. The added salts reduced the toxicity to a minimum, indicating that the lack of essential salts is the primary factor contributing to the increased sensitivity to v-PDMS observed in S-Buffer compared to S-Media (Figure 3a). To further assess the individual contributions of each salt, we tested them separately. As expected, due to its minor contribution to metabolic support, the addition of potassium citrate to S-Buffer did not change mortality.

**Figure 3.**
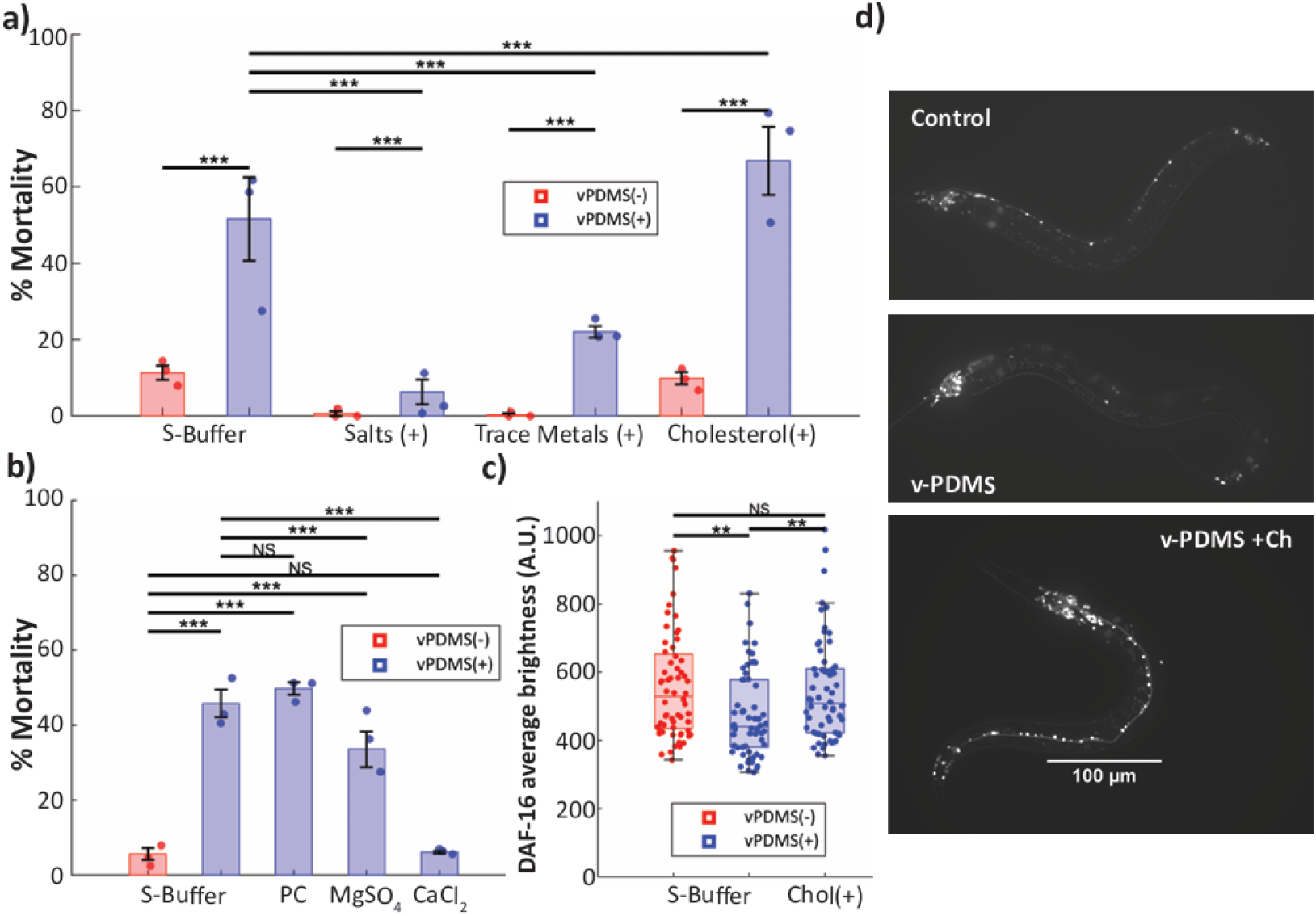
Media components modulate v-PDMS toxicity. (a) Cholesterol and salts supplementation reduces mortality, while trace metals increase it. (b) Addition of Individual salts: potassium citrate (PC) shows no effect, magnesium partially protective, calcium fully protective. Mean ± SEM; Fisher*’*s exact test with Bonferroni correction. (c) DAF-16::GFP expression is reduced after v-PDMS exposure in the absence of cholesterol data analyzed by two-way ANOVA with Bonferroni correction. (d) Example images of DAF-16::GFP expression.

In contrast, magnesium sulfate partially increased the survival rate, while calcium chloride provided complete protection against the toxic effects of v-PDMS (Figure 3b). As described earlier, cholesterol also emerged as a key protective factor. In addition to being crucial for development and reproduction, cholesterol plays a key role in *C. elegans* by enabling the synthesis of sterol-derived hormones required for activation of the DAF-16 stress response pathway^38,39^. We hypothesized that the absence of exogenous cholesterol impairs the activation of a DAF-16 response to v-PDMS exposure. To further investigate how cholesterol confers this protective effect, we used a transgenic *C. elegans* strain expressing fluorescently labeled DAF-16 protein with Green fluorescent protein (GFP), and quantified its abundance under different conditions. We found that worms exposed to v-PDMS without cholesterol supplementation showed a decrease in overall DAF-16::GFP reporter levels compared to control groups with either cholesterol or no v-PDMS exposure (Figure 3c-d). This reduction in reporter signal may reflect altered DAF-16 activity or stability under sterol-limited conditions. However, we did not observe significant differences in nuclear translocation of DAF-16::GFP between these groups (Supplementary Figure 2), suggesting that the observed effect is more likely related to reporter abundance rather than canonical nuclear localization dynamics.

### 2.4. Toxicity of short PDMS oligomers and cyclic siloxanes

PDMS elastomer formulations can include a variety of small linear and cyclic organosilicone compounds, which differ in their potential leachability and bioactivity. Among the short linear siloxanes, formulations may contain hexamethyldisiloxane (MM), octamethyltrisiloxane (MDM), and decamethyltetrasiloxane (MD2M), which vary in chain length and solubility. Additionally, cyclic siloxanes, such as hexamethylcyclotrisiloxane (D3), octamethylcyclotetrasiloxane (D4), and decamethylcyclopentasiloxane (D5), are often present^17^. These compounds are products of the synthesis process^3,4^ or added to serve as processing aids, plasticizers, and have been identified as a cause of cellular death^19^.

To assess the potential biological effects of additional PDMS components, we tested the effects of linear and cyclic siloxanes. The siloxanes were equilibrated in S-Buffer following the same protocol used in previous sections for v-PDMS exposures. Linear siloxanes were tested independently, while the cyclic siloxanes (D3, D4, and D5) were evaluated as a mixture. Cyclic siloxanes were tested as mixtures because D3 is not liquid at room temperature and requires dissolution, making it impractical to test in isolation. Additionally, these cyclic siloxanes are commonly found in a molar ratio of 1:8:1 (D3:D4:D5) in PDMS formulations^17^. By maintaining this ratio, we aimed to replicate the typical composition of cyclic siloxanes present in elastomer formulations and assess their collective impact on *C. elegans*.

The results revealed that the linear siloxanes tested exhibited a statistically significant degree of toxicity compared to the control group, suggesting that these compounds may contribute to the overall negative biological impact of PDMS leachates (Figure 4). In contrast, exposure to the cyclic siloxane mixture (D3:D4:D5, 1:8:1 molar ratio) did not result in any observable difference in survival compared to the control, indicating that these compounds, in the tested conditions, do not contribute to the toxic effects observed in other PDMS components.

**Figure 4.**
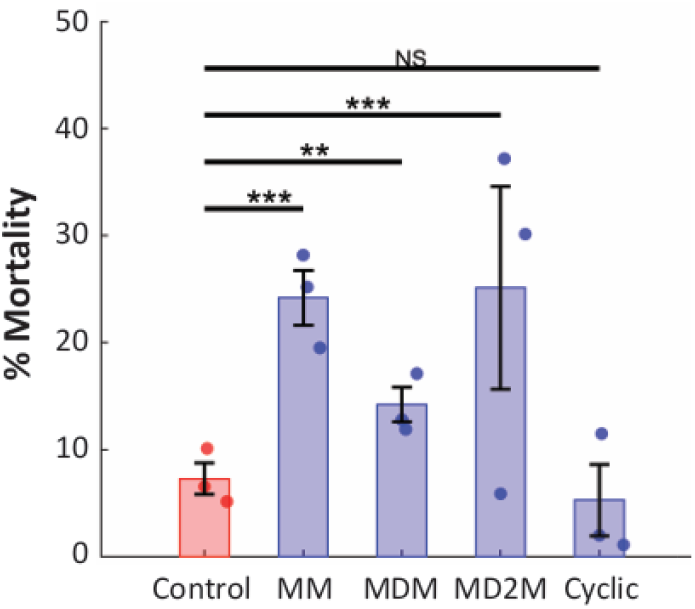
Toxicity of short siloxanes. Exposure to linear siloxanes (MM, MDM, MD2M) slightly but significantly increases mortality, while cyclic siloxanes (D3:D4:D5, 1:8:1) do not differ from controls. Mean ± SEM. Fisher*’*s exact test with Bonferroni correction.

### 2.5. Protective effects of tetrakis-(dimethylsiloxy)-silane crosslinker

Another essential component of a PDMS network is the crosslinker^11,34,40^. Previous studies have reported that the crosslinker tetrakis-(dimethylsiloxy)-silane (TDSS) (Figure 5a) exhibits acute toxicity in other organisms, particularly when combined with v-PDMS oils^17^. To evaluate its potential bioactivity in *C. elegans*, we equilibrated TDSS in S-Buffer using the same protocol described for other PDMS components and exposed L4-stage larvae to the resulting solution. Surprisingly, worms exposed to TDSS alone consistently exhibited lower mortality rates than control groups cultured in plain S-Buffer (Figure 5b), suggesting a possible protective role for the compound under these conditions. Intrigued by this outcome, we proceeded to mix v-PDMS with TDSS (which do not react in the absence of a catalyst) at increasing TDSS concentrations. Remarkably, we observed a dose-dependent protective effect, where higher concentrations of TDSS were associated with increased worm survival, even in the presence of v-PDMS (Figure 5c).

**Figure 5.**
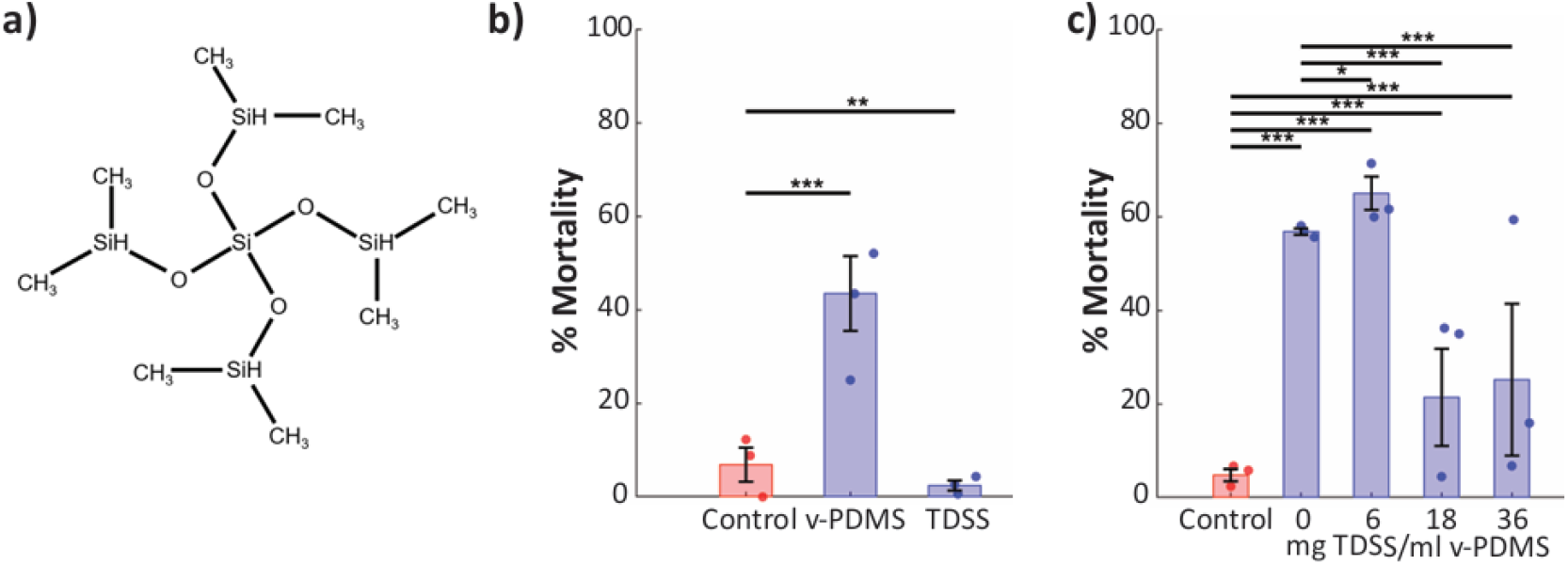
Effects of TDSS crosslinker on *C. elegans* survival. (a) Chemical structure of tetrakis-(dimethylsiloxy)-silane (TDSS). (b) TDSS alone reduces mortality compared to S-Buffer control. (c) TDSS attenuates v-PDMS toxicity in a dose-dependent manner. Mean ± SEM, with star symbols for weighted averages. Fisher*’*s exact test with Bonferroni correction.

These findings suggest that TDSS may counteract or neutralize some of the toxic effects induced by low-molecular-weight v-PDMS chains, though the underlying mechanism remains unclear. One possibility is that TDSS interacts physically or chemically with toxic species in solution, reducing their bioavailability or altering their uptake. Alternatively, TDSS itself could activate a protective response pathway in *C. elegans*, similar to what we observed with calcium chloride and cholesterol, especially given that the TDSS-treated group consistently outperformed the control group in terms of survival. To determine whether TDSS acts through sequestration of toxic species, modulation of their uptake, or induction of a protective response, further mechanistic studies are required.

## 3. Discussion

This study examines the biological effects of PDMS oils on *C. elegans*, specifically investigating how v-PDMS chains and other PDMS-derived compounds influence worm survival under various environmental conditions. Understanding these interactions is important because PDMS is widely used in biomedical applications, microfluidics, and consumer products. Yet, its assumed biocompatibility remains largely untested in whole-organism models, such as *C. elegans*. Our findings reveal that v-PDMS chains that dissolve in aqueous solutions induce mild toxicity in *C. elegans* larvae. When the media provides access to all essential nutrients, larvae do not experience significant detrimental effects from PDMS. However, when cultured in a suboptimal medium, leachates from an undercrosslinked PDMS (5:1) networks can induce death in nematode larvae, suggesting that environmental factors play a critical role in modulating PDMS toxicity.

Essential salts play a crucial role in *C. elegans’* health by maintaining ionic balance, pH stability, and essential physiological functions. In our experiments, the absence of essential salts in S-Buffer resulted in increased sensitivity to v-PDMS, while supplementation with calcium chloride and magnesium sulfate significantly improved survival rates. It is unlikely that the difference in resistance is due to osmotic strength since the majority (>99%) of the ionic strength is provided by the NaCl that is already present in the S-Buffer ^41^, therefore, their protective role is likely to stem from the biological activity of calcium and magnesium. Calcium ions play a critical role in *C. elegans* physiology, particularly in neuronal function, muscle activity, and cellular communication, all of which are necessary for stress resistance. In the nervous system, calcium is a key second messenger that regulates synaptic vesicle release, ensuring efficient neuronal communication ^42^. Proper signaling enables worms to respond to environmental changes, including the presence of toxins, by modulating locomotion, feeding behaviors, avoidance responses, and the activation of defense gene expression programs. Calcium is also essential for muscle contraction, as it facilitates the excitation-contraction coupling mechanism necessary for coordinated movement^43^. Maintaining normal motility is particularly important in toxic environments, where impaired movement can increase vulnerability to stressors. Additionally, calcium plays a broader role in cellular communication by establishing ideal protein folding conditions; a calcium deficit could lead to the misfolding of signaling proteins, compromising cellular homeostasis and the organism*’*s ability to respond to stress^44,45^. Magnesium ions also contribute to stress resilience, but in a different manner. As a vital cofactor for numerous enzymatic reactions, magnesium supports DNA replication, ATP production, and protein synthesis, all of which are essential for maintaining metabolic homeostasis and genomic stability^46,47^. Because metabolic disruption is a common consequence of chemical stress, supplementing with magnesium may help buffer against the harmful effects of PDMS leachates by maintaining cellular processes in a functional state. Additionally, magnesium stabilizes membrane integrity and ion channels^46^, which could help worms regulate osmotic balance and maintain normal physiological function in varying environmental conditions.

We observed that, under the availability of cholesterol, *C. elegans* larvae acquire some resistance to the v-PDMS. Cholesterol, even in small concentrations, is indispensable for the development of *C. elegans* into adulthood and reproduction since they cannot synthesize sterols de novo^38,48^. Sterol-derived hormones play a crucial role in regulating genes in the DAF family^48,49^. Cholesterol-dependent resistance to v-PDMS may result from hormonal activation of DAF-16, a master regulator of longevity and starvation stress response in *C. elegans*. DAF-16 functions downstream of DAF-12, a nuclear receptor that requires cholesterol-derived hormones for activation^38^. In our experiments, worms exposed to v-PDMS without cholesterol supplementation exhibited significantly lower DAF-16 activation compared to worms exposed to v-PDMS with cholesterol and to those cultured without either v-PDMS or cholesterol supplementation. These findings suggest that cholesterol in the medium may play a protective role by enabling the biosynthesis of sterol-derived hormones necessary to trigger the DAF-12/DAF-16 mediated stress response pathway. The transgenerational resistance observed in worms whose ancestors experienced early-life starvation supports the hypothesis that the DAF-16 pathway may play a role in protection against PDMS toxicity. The DAF-16 gene is a key regulator of development and stress response, and is known to be activated under conditions of reduced insulin signaling and dietary restriction to enhance survival during starvation^50^. The absence of cholesterol in the medium could impair the worms*’* ability to properly activate these protective pathways, thereby increasing their susceptibility to PDMS toxicity. This effect may be further exacerbated by the sequestration of sterol-derived hormones by PDMS present in the medium. Previous studies have reported that PDMS can adsorb hydrophobic sterol-like molecules when cells are cultured on its surface^13^. Given that sterol-derived hormones are present in *C. elegans* at inherently low concentrations, even a small degree of sequestration could be sufficient to disrupt critical biological processes, potentially compounding the effects of PDMS toxicity. Another possible mechanism is that cholesterol deprivation alters membrane integrity and permeability^51^, facilitating the entry of hydrophobic PDMS molecules into cells and thereby amplifying their toxic effects. We did not directly test this possibility in our study. Further work is required to determine whether altered membrane properties contribute to the increased mortality observed under cholesterol starvation.

Trace metals are essential for cellular function, playing key roles in signaling pathways, protein synthesis, and enzymatic activity^52^. However, at elevated concentrations, they can be toxic to organisms, and their ability to clear them is limited due to poor diffusion across membranes^53^. Instead, organisms rely on specialized transporters to regulate intracellular metal levels and maintain homeostasis^52^. In our assays, trace metals alone did not induce mortality in *C. elegans*. However, when worms were exposed to v-PDMS oils in the presence of trace metals, we observed a significant increase in lethality. Under standard culture conditions, worms are typically exposed to trace metals only in the presence of bacterial food, allowing the metals to be first processed by *E. coli* and later assimilated indirectly through ingestion. In contrast, our experimental setup lacked food, and thus, worms were directly exposed to elevated concentrations of free trace metals in the medium. This may have compounded physiological stress and increased susceptibility to v-PDMS-induced toxicity. Interestingly, when worms were cultured in v-PDMS-equilibrated S-Medium, which contains the same concentration of trace metals as in the supplemented S-Buffer assays, no significant increase in mortality was observed. This suggests that the protective components of S-Medium (calcium chloride, magnesium sulfate, and cholesterol) are sufficient to counteract the negative effects of both trace metals and v-PDMS. The presence of calcium and magnesium likely contributes to this protection by supporting cellular homeostasis, stabilizing membranes, and competitively inhibiting the uptake of toxic metal ions. These ions also play critical roles in maintaining ion channel function and activating stress response pathways, which may collectively enhance the worms’ resilience to chemical stressors.

Aside from the functionalized v-PDMS, we identified that short PDMS molecules also exhibit mild toxicity in *C. elegans*. Our findings indicate that linear siloxanes, such as hexamethyldisiloxane, octamethyltrisiloxane, and decamethyltetrasiloxane, induced a small but statistically significant increase in lethality, suggesting that these compounds may contribute to the negative biological effects of PDMS leachates. Cyclic siloxanes (D3, D4, D5) have been shown to induce cell death in human cell lines^19^. However, when we tested their effects on *C. elegans*, no significant toxicity was observed. This suggests that their interaction with whole organisms may be more limited under the conditions tested. Moreover, while some PDMS-derived molecules can impact worm survival, their effects depend on chemical structure, environmental solubility, and potential biological interactions. Further studies should investigate whether these effects extend beyond acute lethality, such as metabolic disruptions or sub-lethal physiological impairments, particularly in prolonged exposures.

The protective effect of TDSS was unexpected, and the mechanisms underlying this phenomenon remain unclear. While previous studies have reported the acute toxicity of TDSS in other organisms, particularly when combined with v-PDMS (Mw=17Kda)^17^, our results show that TDSS alone is not toxic to *C. elegans* but also appears to enhance survival. Furthermore, when mixed with v-PDMS, TDSS significantly reduced the lethality of v-PDMS leachates in a dose-dependent manner. The specific biochemical or physical mechanisms responsible for this protective effect are beyond the scope of the current study, but they warrant further investigation. Notably, we could not find any previous studies reporting a protective effect of TDSS at the whole-organism level, making this finding particularly intriguing. Future research should explore whether TDSS influences PDMS bioavailability, adsorption properties, or interactions with biological membranes, which could help explain the observed increase in survival.

Overall, our study provides new insights into the biological interactions between *C. elegans* and PDMS-derived compounds, revealing that while v-PDMS and certain linear siloxanes exhibit mild toxicity, their effects are modulated by environmental conditions and access to essential nutrients. Perhaps most unexpectedly, TDSS displayed a protective effect against v-PDMS toxicity, highlighting the complexity of PDMS-related interactions in biological systems. Given the widespread use of PDMS in biomedical and consumer applications, these results underscore the need for further research into the long-term effects of PDMS exposure and the mechanisms underlying its interactions with biological systems. Future studies should investigate whether these findings extend to other organisms, explore potential metabolic disruptions beyond acute toxicity, and determine the broader implications of PDMS leachates in real-world exposure scenarios.

## 4. Materials and Methods

### Preparation of leachate solutions

Three distinct PDMS elastomer networks were prepared using DOW Sylgard-184 by mixing the elastomer base (Part A) and curing agent (Part B) at weight ratios of 5:1, 10:1, and 20:1 (A:B). Mixtures were vigorously stirred and subsequently degassed in a vacuum chamber for approximately 30 minutes at room temperature. The degassed mixtures were then poured into polystyrene Petri dishes and cured at 65⍰°C for 4 hours, following the manufacturer*’*s recommendations.

After curing, the solid PDMS networks were allowed to return to room temperature and then mechanically crushed using a stainless-steel spice grinder. Ten grams of each cured formulation were transferred into 15⍰mL conical centrifuge tubes, to which 5⍰mL of S-Buffer was added. The mixtures were allowed to equilibrate at room temperature for 24 hours. Following incubation, the liquid was separated by filtering through a Nylon Corning cell strainer with 70um pores into clean tubes and used immediately for subsequent toxicity assays.

Fragments of the shredded PDMS networks were measured by spreading them on the scanning surface of an Epson perfection V600 scanner at maximum resolution. The images were processed with ImageJ by adjusting eliminating the noise with a Band pass filter and adjusting contrast and brightness to obtain individual and full fragments. The scale was set by scanning a 3 inch glass slide alongside the fragments. Then we used the *“*analyze particles*”* function of ImageJ to obtain a list of the areas of the particles with a minimum size of 0.01 in^2^ as it was the threshold we identified to work best to discriminate true particles from background noise (Supplementary Figure 3).

### Preparation of oil mixture solutions

All tested oil-based substances were prepared using a standardized equilibration protocol. In 15⍰mL conical centrifuge tubes, the aqueous medium (S-Buffer or S-Medium) and the oil phase were combined at a 10:1 volume ratio (aqueous:oil). For mixed formulations such as TDSS/v-PDMS and the cyclic siloxane blend, the oils were first pre-mixed by gentle stirring prior to addition to the aqueous phase. All mixtures were then vigorously vortexed and sonicated for 3 minutes to enhance oil dispersion.

This procedure was applied identically to all experimental and control samples. Following mixing, the tubes were left undisturbed at room temperature for 24 hours to allow for phase separation. After incubation, the aqueous phase was carefully transferred to a new centrifuge tube, taking care to avoid any carryover of undissolved oil droplets. The resulting equilibrated media were used immediately for the assays.

### Nematode culture and growth

For all survival experiments, we used *C. elegans* worms from the Bristol N2 wild-type strain. Worms were age-synchronized using a standard bleaching solution (2 parts 1M NaOH, 1 part sodium hypochlorite, 1 part deionized water) and cultured on a nematode growth medium (NGM) agar substrate seeded with *E. coli* OP50 at 20°C for approximately 50 hours, or until they reached the L4 stage^54^. To prevent the activation of epigenetic stress-response pathways, all worms used in these experiments were maintained well-fed for at least five generations unless specified otherwise. Some strains were provided by the CGC, which is funded by NIH Office of Research Infrastructure Programs (P40 OD010440).

### Toxicity assays

Once worms reached the L4 stage, they were washed off the plate with M9 buffer containing 1% Triton X-100 into a 1 ml conical tube, allowed to settle, and then transferred into a 24-well plate by pipetting 5ul from the bottom of the tube. Each well contained 300 µL of either liquid media or the leachate solution of interest. Each experiment included approximately 100 worms per biological replicate, distributed across three wells, with three biological replicates per experiment. Worms were incubated on a rocker at 30 rpm to maintain constant agitation for 24 hours, after which survival was assessed. Worms that failed to exhibit any movement in response to a gentle touch stimulus were recorded as dead.

### DAF-16 imaging

For the DAF-16 expression experiments, we used the ASM10 strain (*daf-16*(del2[*daf-16::GFP-C1^3xFlag*])). Worms were cultured on NGM plates until the L4 stage and then transferred to liquid suspension in 24-well plates 6 hours prior to imaging. After exposure, worms were collected into microcentrifuge tubes, allowed to settle, and transferred using a micropipette. A drop of worms was placed on 2% agarose pads on glass slides, and immobilization was achieved by adding a drop of 2 mM tetramisole in M9 buffer.

Images were collected on a Leica DMi8 microscope fitted with a CrestOptics X-light V2 spinning disk unit and a Hamamatsu Orca-Fusion camera, using a 63× objective lens. Illumination was provided by an 89 North LDI laser diode system. Acquisition parameters were kept identical across all samples: 60ms exposure and 30% laser power. Image stacks consisted of 20 optical sections acquired at 1-µm intervals, and maximum-intensity projections of these z-stacks were used for analysis.

### Chemicals

α,ω-terminated polydimethylsiloxanes with molecular weights 6.0, 28.0, 49.5, and 62.0 kDa (v-PDMS), tetrakisdimethylsiloxysilane (TDSS), hexamethyldisiloxane (MM), octamethyltrisiloxane (MDM), decamethyltetrasiloxane (MD2M), hexamethylcyclotrisiloxane (D3), octamethylcyclotetrasiloxane (D4), and decamethylcyclopentasiloxane (D5) were purchased from Gelest Inc., and used as received.

