## Supplementary figures and images for "Assessing PDMS Biocompatibility in Microfluidic Applications: Toxicity and Survival Outcomes in *C. elegans*"

### Supplementary Figure 1

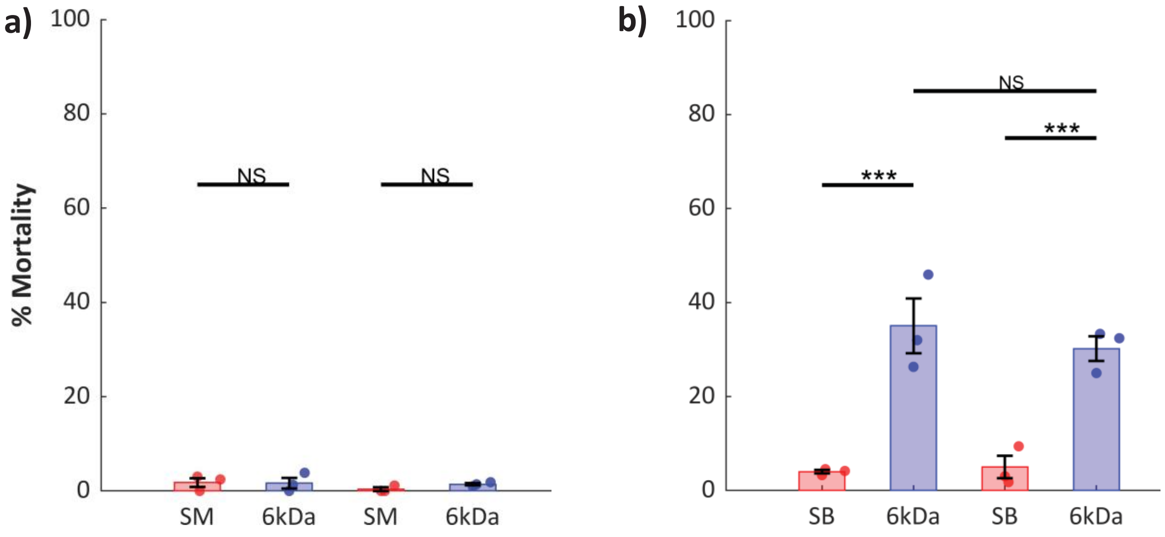

### Supplementary Figure 2

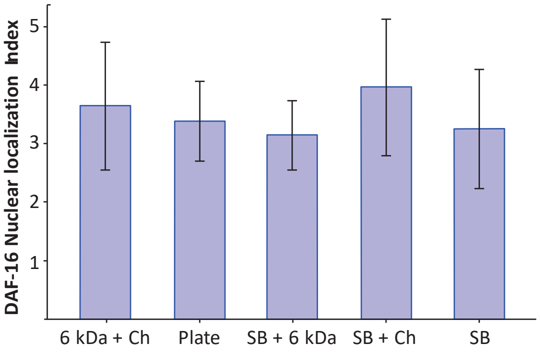

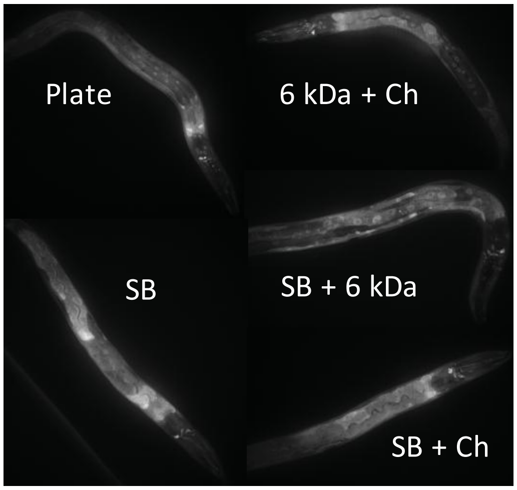

### Supplementary Figure 3

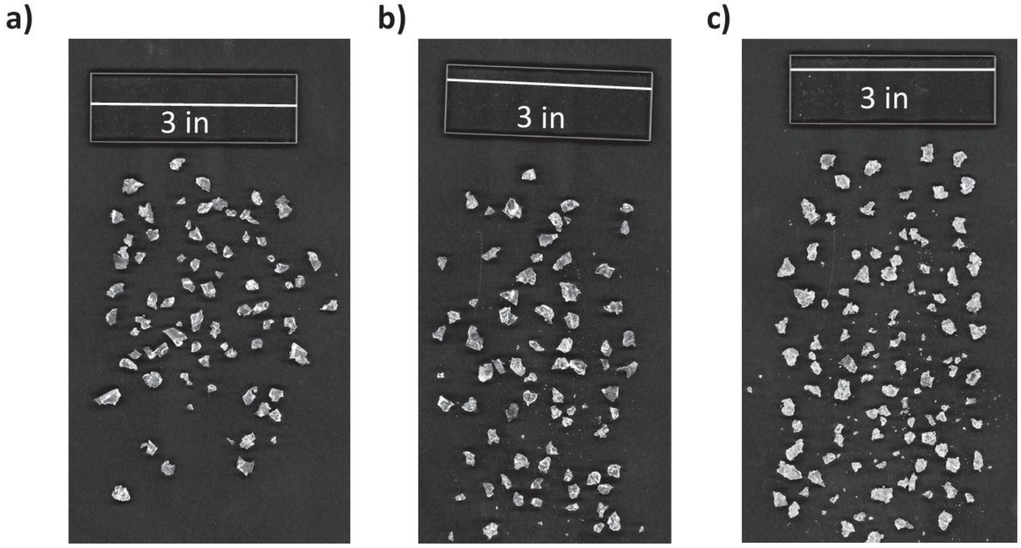
