## Supplementary Table 1 for "Assessing PDMS Biocompatibility in Microfluidic Applications: Toxicity and Survival Outcomes in *C. elegans*"

| Measurement | S-Buffer | S-Media | Salts(+) | Trace Metals (+) | Cholesterol (+) | PC(+) | MgSO4 | CaCl2 |
|---|---|---|---|---|---|---|---|---|
| 1 | 5.91 | 5.80 | 5.81 | 5.89 | 5.88 | 5.80 | 5.88 | 5.89 |
| 2 | 5.97 | 5.81 | 5.80 | 5.89 | 5.89 | 5.80 | 5.90 | 5.91 |
| 3 | 5.90 | 5.80 | 5.79 | 5.90 | 5.88 | 5.80 | 5.88 | 5.88 |
